# Use of an automated pyrosequencing technique for confirmation of Sickle Cell Disease

**DOI:** 10.1101/610063

**Authors:** CC Martino, CS Alencar, P Loureiro, AB Carneiro-Proietti, CA Máximo, RA Mota, DOW Rodrigues, N Gaburo, S Kelly, EC Sabino, for the International Component of the NHLBI Recipient Epidemiology and Donor Evaluation Study (REDS-III)

**Author notes:** **Corresponding author:** Ester C. Sabino.

## Abstract

**Background:** The diagnosis of sickle cell disease (SCD) is made by hemoglobin assays such as high-performance liquid chromatography (HPLC), isoelectric focusing and cellulose acetate or citrate agar electrophoresis. These assays are easy to perform and used in large-scale newborn screening in many countries. These tests however may not easily differentiate Sβ^0^ thalassemia from SS or identify other hemoglobin variants, and in this case, hemoglobin (HBB) gene sequencing may be necessary.

**Objectives:** To develop a high throughput DNA based confirmatory assay for SCD and to detect mutations in the HBB gene.

**Methods:** We developed an automated pyrosequencing technique (PyS) based on QIAGEN technology (Hilden, Germany) to detect homozygous or heterozygous hemoglobin S mutations as well as hemoglobin C mutations. The technique was tested on 2,748 samples from patients enrolled in a multi-center SCD cohort in Brazil. Patients were previously tested using HPLC to diagnose SCD as part of routine clinical care. Any subjects with discrepant results between HPLC and PyS or with heterozygous hemoglobin S detected had Sanger sequencing of the HBB gene.

**Results:** We identified 168 samples with discrepant results between HPLC and PyS and 100 with concordant HPLC and PyS= heterozygous S, which would suggest Sβ-thalassemia or other hemoglobin S variants. The PyS assay correctly identified 1906 (98.7%) of the 1930 HbSS and 628 (98.7%) of the 636 HbSC samples. Of the 179 remaining samples, PyS correctly indicated S heterozygosis in 165 (92.2%). Of the 165 heterozygous S samples confirmed by Sanger as consistent with Sβ thalassemia genotype, 84 samples were classified as Sβ^0^ thalassemia and 81 as Sβ^+^ thalassemia. The most frequent beta thalassemia mutations of Sβ^0^ and Sβ^+^ were HBB: c.118C>T (Gln40Stop) and HBB c.92 + 6T> C, respectively.

**Discussion:** The PyS proved to be satisfactory for large-scale confirmatory testing of hemoglobin mutation. Moreover, with this study we were able to describe the most common β^+^ and β^0^ mutations in SCD patients with Sβ-thalassemia in a large multi-institutional SCD cohort in Brazil.

## INTRODUCTION

Sickle cell disease (SCD) is an inherited red blood cell disorder in which at least one of the HBB genes has a Glu6Val mutation. When both genes are mutated (SS) the individual demonstrates a severe form of the disease typically called sickle cell anemia (SCA). The coinheritance of HbS with other abnormal β-globin chain variants can also cause SCD (1–3).The most common mutations are sickle-hemoglobin C disease (HbSC) and sickle β-thalassemia (Sβ^+^ thalassemia and Sβ^0^ thalassemia). Mutations designated as β0-thalassemia are associated with no normal hemoglobin A production, therefore clinical symptoms of Sβ^0^ thalassemia are typically as severe as SS and also usually classified as sickle cell anemia (4–6). Mutations designated as β^+^-thalassemia are associated with variable levels of normal hemoglobin A.

The diagnosis of SCD is made by hemoglobin assays such as high-performance liquid chromatography (HPLC), isoelectric focusing, cellulose acetate electrophoresis and citrate agar electrophoresis. Those assays are easy to perform and used in large scale newborn screening in many countries including Brazil. The tests however may not easily differentiate Sβ^0^ thalassemia from SS, and in this case HBB gene sequencing is necessary (7,8).

In 2013, a large multi-center cohort was established in Brazil to characterize clinical outcomes in the Brazilian SCD population under the National Heart Lung and Blood Institute Recipient Epidemiology and Donor Evaluation Study -III (REDS-III) program (9). The genotype of the participants was defined by each site was based on HPLC measurement of variant hemoglobins, however the results were classified differently by each site. A genotype confirmation based on DNA was necessary to ensure standardized classification of SCD genotype for the research.

Because the HbS and HbC mutations are separated by only one nucleotide, it is not easy to develop specific probes for real time PCR (10,11). We describe here a pyrosequencing technique (PyS) that was developed to confirm the SCD genotype for participants in the REDS-III Brazil SCD study. The technique was validated using Sanger sequencing of the HBB gene as the gold standard. This approach also allowed us to describe the most common HBB mutations in patients classified as Sβ^0^ thalassemia and Sβ^+^ thalassemia.

## MATERIALS AND METHODS

### Samples

This study was performed using samples collected for the REDS-III Brazil SCD cohort study (9) eligible participants were randomly selected that included institutions in four Brazilian states: São Paulo (Hospital das Clínicas), Minas Gerais (Hemominas), Rio de Janeiro (Hemorio) and Pernambuco (Hemope). The samples were collected in an EDTA tube, centrifuged at 3500rpm, and plasma was separated from cells. Both components were frozen and shipped to the central laboratory at the University of Sao Paulo for further testing.

DNA extraction was performed using the QIAsymphony apparatus (Qiagen, Germany) and the QIAsymphony DNA Mini Kit (Qiagen, Germany), following the manufacturer’s instructions and protocol.

### Pyrosequencing

Primers sequences were designed using the Pyromark Assay by Design software as follows: Forward 5’ATTGCTTACATTTGCTTCTGACAC3’, Reverse 5’ACCAACTTCATCCACGTTCAC3’, targeting the same regions proposed by Sutton, Bouhassira (12). PCR was performed with the PyroMark PCR Kit (QIAGEN) using 100 ng / μL of DNA according to the manufacturer’s protocol. The PCR product was used for the pyrosequencing assay with PyroMark Q24 Gold Kit (Qiagen, Germany) and subsequently subjected to PyroMark Q24 sequencer (Qiagen, Germany) using the primer sequence 5’CATGGTGCATCTGACT3’. The analysis was performed using Pyrogram (PQ24 Software) version 2.1 (Qiagen, Germany).

### Sanger sequencing

We used the Sanger sequencing technique for confirmation of discrepant samples and to make the determination of β thalassemia mutations. PCR was used to amplify a fragment of 101 base pairs covering the coding region of the Beta Globin gene, which has approximately 619 base pairs, using the follow primers sequence: (P1) 5’-TCCTAAGCCAGTGCCAGAAG-3’ and the downstream primer (P5) 5’-TCATTCGTCTGTTTCCCATTC3’(13).

The purified PCR product was subjected to another PCR reaction using the ABI PRISM Big Dye Terminator Cycle Sequencing Ready Reaction kit (Applied Biosystems Foster City, CA) following the manufacturer’s protocol. Subsequently, the products of this reaction were analyzed by an ABI3500 automated sequencer (Applied Biosystems).

The sequences were edited through Sequencher Software (GENECODES) and the results were classified as β^+^ or β^0^ thalassemia mutations previously described in the literature through an online tool HbVarDatabase, Inc. (http://globin.bx.psu.edu/hbvar/) (14,15).

### TOPMed

After establishment of the cohort, the REDS-III Brazilian SCD cohort was selected to participate in the National Heart Lung and Blood Institute Trans-Omics for Precision Medicine (TOPMed) Program, which generates whole-genome sequencing and other -omics data on well phenotyped cohorts. The program will integrate -omics data with molecular, behavioral, imaging, environmental, and clinical data to improve the prevention and treatment of blood and other disorders (16).

Whole genome sequencing was performed in samples of the REDS-III Brazilian SCD cohort by sequencing centers to a median depth of 39X using DNA from blood, PCR-free library construction and Illumina HiSeq X technology (nhlbiwgs.org). These sequences were utilized in the present research as a means of final confirmation of the SCD genotype in combination with Sanger sequences. Results of pyrosequencing assay were compared with final SCD genotype classification.

## RESULTS

The REDS-III Brazilian SCD cohort study enrolled 2,793 patients from 2013-2015. A total of 2,749 samples were obtained from the first visit of the enrolled patients. The number of patients per site classified by their original HPLC results is summarized in Table 1. The center from Rio de Janeiro (Hemorio) combined SS and Sβ^0^ thalassemia in the same category, while the centers from Minas Gerais (Hemominas) combined Sβ^0^ and Sβ^+^. São Paulo and Pernambuco provided results that classified patients as SS, Sβ^0^ and Sβ^+^ separately.

**Table 1.**
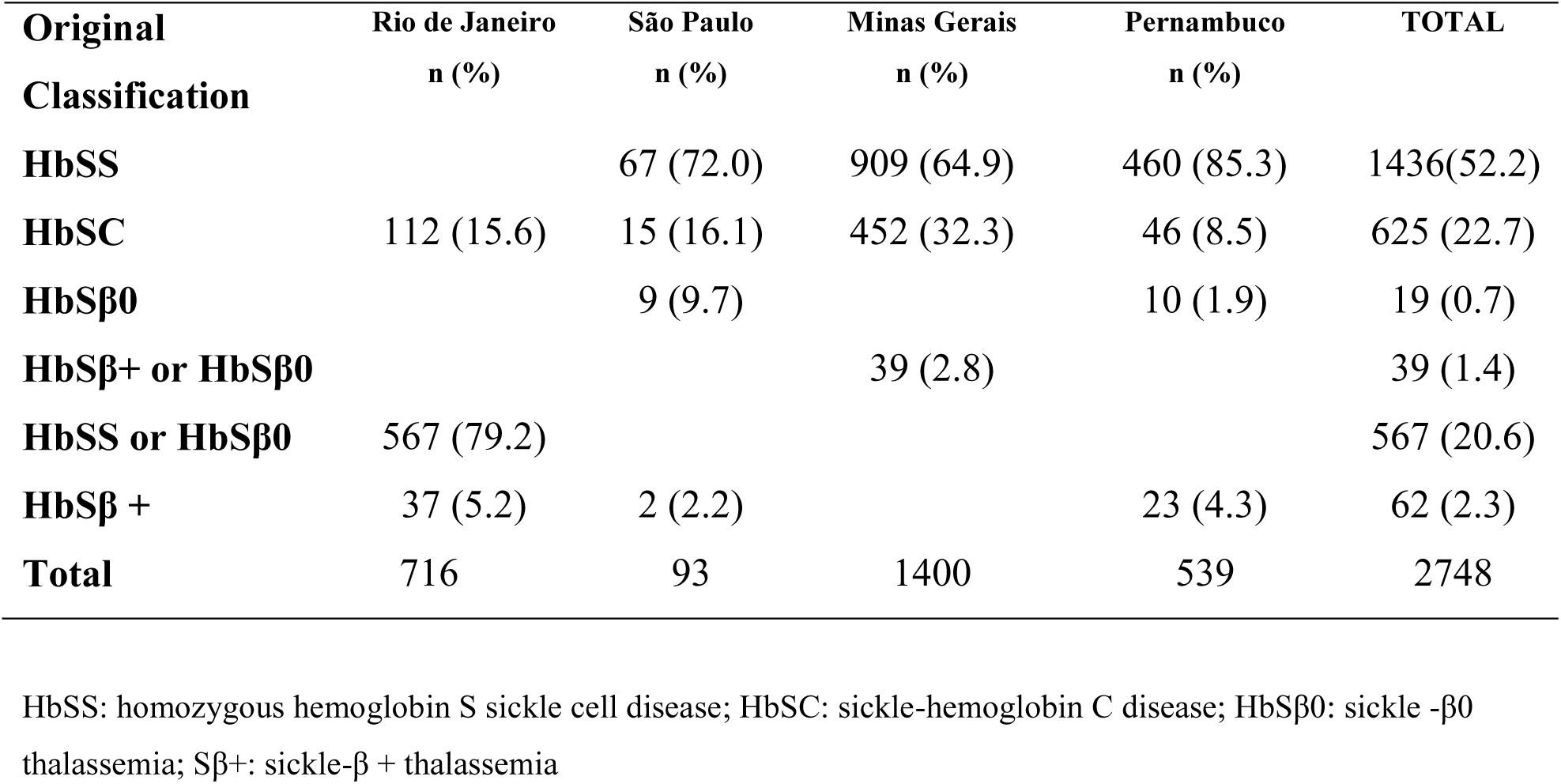
Classification of SCD genotype of REDS-III Brazil SCD cohort participants provided by each center using high-performance liquid chromatography

All samples with heterozygous S (n=100) results or discrepant results between HPLC and PyS (n=168) were submitted to Sanger sequencing of the HBB gene to assign a REDS-III SCD genotype classification. When the TOPMed whole genome sequencing later became available, we compared all the results. Twenty samples with discrepant results between REDS-III final classification and TOPMed classification were repeated using Sanger sequence and a final genotype was then assigned. Comparison of the PyS results with the final confirmed classification of SCD genotype is shown in Table 2. The PyS assay correctly identified 1906 (98.7%) of the 1930 HbSS and 628 (98.7%) of the 636 HbSC samples. Of the 179 remaining samples, PyS correctly indicated S heterozygosis in 165 (92.2%).

**Table 2.** Comparison of pyrosequencing results with final SCD genotype classification within the REDS-III Brazil SCD cohort study

| Final SCD Classification | Pyrosequencing |  |  | TOTAL |
| --- | --- | --- | --- | --- |
|  | HbSS | HbSC | Hb S |  |
| **HbSS** | **1903** | 12 | 15 | 1930 |
| **HbSC** | 8 | **628** |  | 636 |
| **HbSβ+** | 9 |  | **72** | 81 |
| **HbSβ<sup>0</sup>** | 4 |  | **80** | 84 |
| **HbSD** | 1 |  | **6** | 7 |
| **HbS/HPFH** | 3 |  |  | 3 |
| **HbSJ** |  |  | **2** | 2 |
| **S/Quebec-Chori** |  |  | **1** | 1 |
| **S/K-Woolwich** |  |  | **1** | 1 |
| **S/Korle-Bu** |  |  | **1** | 1 |
| **S/Porto Alegre** |  |  | **1** | 1 |
| **S/Deer Lodge** |  |  | **1** | 1 |
| **TOTAL** | 1934 | 640 | 180 | 2748 |

HbSS: homozygous hemoglobin S sickle cell disease; HbSC: sickle-hemoglobin C disease; HbSβ<sup>0</sup>: sickle -β<sup>0</sup> thalassemia; Sβ+: sickle-β + thalassemia; Hb S: heterozygous S HPFH: hereditary persistence of fetal hemoglobin; Samples correctly identified by pyrosequencing are shown in bold

Sanger sequencing allowed us to define the beta thalassemia mutations in the study population. The distribution of mutations varied according to the regions studied (table 3). The most common mutation was a β^0^ mutation, HBB: c.118CT(Gln40Stop) [codon 39 (C>T)], in all sites with exception of Pernambuco, where the β^+^ mutation HBB:c.92+5G>C [IVS-I-5 (G>C)] was more common. In the state of Rio de Janeiro we identified one rare HBB mutation: HBB:c.75T>A [codon 24 (T>A)], a variant that leads to mild Sβ+ thalassemia.

**Table 3.** Classification of beta thalassemia mutations, REDS-III Brazil SCD cohort study.

| Mutations | β-Thal. | MG | PE | RJ | SP | n (%) |
| --- | --- | --- | --- | --- | --- | --- |
| HBB:c.118C>T(Gln40Stop)[codon 39 (C>T)] | Sβ0 | 26(38.8) | 6(14.0) | 10(23.3) | 5(41.7) | 47 (28.5) |
| HBB:c.92+6T>C [IVS- I-6 (T>C)] | Sβ+ | 6(8.9) | 10(23.3) | 7(16.3) | 2(16.7) | 25 (15.2) |
| HBB:c.92+1G>A [IVS-I-1 (G>A)] | Sβ0 | 11(16.4) | 4(9.3) | 5(11.6) | 1(8.3) | 21 (12.7) |
| HBB:c.93-21G>A [IVS-I-110 (G>A)] | Sβ+ | 7(10.4) | 3(7.0) | 5(11.6) | 2(16.7) | 17 (10.3) |
| HBB:c.92+5G>C [IVS-I-5 (G>C)] | Sβ+ | 1(1.5) | 11(25.6) | 2(4.7) | 1(8.3) | 15 (9.1) |
| HBB:c.-79A>G(-29AG) [(-29A>G)] | Sβ+ | 3(4.5) | | 6(14) | | 9 (5.5) |
| HBB:c.92+5G>A [IVS-I-5 (G>A)] | Sβ+ | 2(3.0) | 1(2.3) | 3(7) | 1(8.3) | 7 (4.2) |
| HBB:c.315+1G>A [IVS-II-1 (G>A)] | Sβ0 | 4(6.0) | | 2(4.6) | | 6 (3.6) |
| Htz IVS-II-849 (A>G) | Sβ0 | 1(1.5) | 2(4.6) | | | 3 (1.8) |
| Htz PolyA, AATAAA>AACAAA | Sβ+ | 2(3.0) | | | | 2 (1.2) |
| Htz -88 (C>T) | Sβ+ | | 2(4.6) | | | 2 (1.2) |
| HBB:c.-138C>T | Sβ+ | | 1(2.3) | | | 1 (0.6) |
| Htz IVSII-839(T>C) Htz IVSII-844 (C>A) | Sβ+ | | | 1(2.3) | | 1 (0.6) |
| DELEÇÃO 572het_deIG | Sβ0 | 1(1.5) | | | | 1 (0.6) |
| HBB:c.321_322insG Htz Cod 106/107(+G) | Sβ0 | | 1(2.3) | | | 1 (0.6) |
| HBB:c.92+2T>C Htz IVSI-2 (T>C) | Sβ0 | 1(1.5) | | | | 1 (0.6) |
| HBB:c.75T>TA | Sβ+ | | | 1(2.3) | | 1 (0.6) |
| p.Glu7Glyfs | Sβ0 | | | 1(2.3) | | 1 (0.6) |
| HBB:c.92+2T>G | Sβ0 | | 1(2.3) | | | 1 (0.6) |
| Htz Stop+4 (C>T) | Sβ+ | 1(1.5) | | | | 1 (0.6) |
| No mutation found in the exon 1 and 2* | Sβ+/ Sβ0 | 1(1.5) | 1 (2.3) |  |  | 2 (1.8) |
| TOTAL |  | 67 | 43 | 43 | 12 | 165 |

\* No DNA available for further sequences

An overall summary of the original classification made by HPLC at the participating sites, samples submitted to Sanger sequencing and final REDS-III SCD genotype classification considering our results compared to whole genome sequencing generated by TOPMed is shown in Figure 1.

## DISCUSSION

There is a need for rapid and precise methods to facilitate the diagnosis of hemoglobinopathies, especially in situations in which conventional testing may not be possible or reliable. For example only frozen samples, in which hemoglobin based assays are less reliable, were available for this research study. The ability to differentiate SS from Sβ0 thalassemia is also not always possible using hemoglobin based assays as nearly all hemoglobin detected is hemoglobin S with no hemoglobin A present. In the absence of information regarding hemoglobin mutations in parents or other clinical and laboratory testing, DNA based testing is required to confirm the SCD genotype. However, the vast majority of cases would be expected to be homozygous SS and sequencing a large number of samples to separate the two would be labor intensive and cost prohibitive. Pyrosequencing is relatively quick and simple and also allows a large scale approach to provide timely diagnosis. In the present study we used the pyrosequencing technique to classify the hemoglobinopathy diagnosis of participants in a large multi-institutional cohort study of SCD by confirming HbSS, HbSC and heterozygous S participants. This allowed targeted Sanger sequencing only in participants with results not concordant with clinical diagnosis assigned at treating center (n=165) and in heterozygous S samples (n= 100) for identification of hemoglobin mutations. The pyrosequencing assay correctly identified 2,699 (98.2%) of the samples and proved to be a satisfactory technique for large-scale testing.

The pyrosequencing technique is a highly reliable tool for the determination of small regions inside the globin genes, and has the advantage of being a relatively simple technique. In addition, pyrosequencing is faster and is associated with a lower cost of operation when compared to other sequencing methodologies. (17). There were 49 samples misclassified by PyS, mostly due to the low level of the pyrogram peaks, which could be improved by standardizing the peak levels below which the batch should be repeated.

In this study we also described the most common β^+^ and β^0^ thalassemia mutations among Sβ thalassemia cohort participants in four states of Brazil. Of the heterozygous S samples confirmed by Sanger, 84 were classified as Sβ^0^ thalassemia and 81 as Sβ^+^ thalassemia.

The types of beta thalassemia mutations demonstrated in this cohort reflect the genetic diversity of the study population. The Brazilian population is the result of admixture between different groups at different time periods; the colonizing Spaniards mixed with the indigenous populations as well as with African slaves during three centuries. Later, immigration from Spain, Italy, Portugal contributed further to the admixture of the present-day Brazilians(18).

The most frequent mutation in our subjects, HBB:c.118C>T(Gln40Stop) Codon 39 (C>T), is also the most prevalent Sβ thalassemia mutation in the Mediterranean. It is believed that codon 39 (C> T) is of Roman origin, and has a high prevalence in Sardinia, mainland Italy, Spain, Portugal and Tunisia (19). Different studies also found this mutation to be frequent in Venezuela (18), Northern Greece (20), Syria (21), and confirmed it in Tunisia (22) and Italy (23).

The next most common β0 thalassemia mutation in our cohort, IVS-I-1, shows a restricted geographical distribution in Eastern Mediterranean countries (Syria, Lebanon, Jordan, Palestine and Egypt) (21).

The presence of the IVS-I-6 mutation, the most common β+ mutation in our cohort, appears to be a contribution from the Portuguese to the genetic makeup of the population, as it corresponds to 29.4% of the alleles in β+ mutations in Portugal. (18,24).

Our results are in accordance with previous Brazilian studies (24–27). As expected, considering the migratory activity of the Brazilian population and ethnic ancestry, the pattern observed is similar to the Mediterranean populations. Interestingly, a study identifying mutations in 31 Sβ thalassemia patients in the state of Rio Grande do Norte did not identify the mutation Codon 39 (C>T) that was common in our study and others in Brazil (28). They identified 15 (48.4%) patients with the IVS-I-1 mutation, 13 (41.9%) with the IVS-I-6 mutation, 2 (6.5%) with the IVS-I-110 mutation and 1 (3.2%) with IVS-I-5 mutation.

Different from the other states in our study, the most common mutation in the state of Pernambuco was IVS-I-5 (G>C). This mutation is very common in Asia, especially in Malaysia and Indonesia and in several regions of India (29). In studies conducted by Khan et al. from 2011-2013 in four provinces of Pakistan, the most frequent mutation detected in a total of 63 samples of β-thalassemia was IVS-I-5(G>C) (33.9%) (30). In India, more than 90% of mutations in beta thalassemia involve IVS1-5 (G >C) (31,32). Similar to our findings, studies by Silva (33) and Araujo (34) also identified this mutation in the population of Recife, Pernambuco. In the 17th century Recife was an important commercial harbor, it is possible that people from the Indian subcontinent (Goa) were brought as slaves by the Portuguese to that area (34).

The racial heterogeneity of the immigrant population in a non-endemic country significantly increases the spectrum of hemoglobinopathy mutations and their combinations found in individuals, making the provision of a molecular diagnostic prenatal diagnosis service more challenging. With the testing algorithm described, it was possible to determine the spectrum of Sβ thalassemia mutations and their combinations in a Brazilian SCD population. It is important to determine the correct mutations for genetic counseling and to identify patients potentially eligible for new drugs or gene therapy trials that may be available for targeted populations. (35).

In conclusion, the pyrosequencing technique is a highly reliable tool for the classification of SCD and is suitable for large-scale testing to identify hemoglobin S (homozygous or heterozygous carriers) and C mutations. This allows targeted hemoglobin sequencing in a limited number of patients, facilitating proper diagnosis when conventional techniques may have limited ability and ensuring proper hemoglobinopathy diagnosis which is essential for correct screening and treatment strategies for patients with SCD.

## ACKNOWLEDGEMENTS

The authors acknowledge all of the research staff at the participating blood centers in Brazil who have enrolled patients into the study and completed all study procedures. At each site the following specific people are recognized for their commitment and contribution to this project: Fundação Pró-Sangue (São Paulo) – Alfredo Mendrone Jr., Cesar de Almeida Neto; ITACI – Instituto de Tratamento do Câncer Infantil (São Paulo) – Roberta Carlucci, Erivanda Bezerra; Hemominas – Belo Horizonte (Minas Gerais) – Carolina Miranda, Tassila Salomon, Franciane Mendes de Oliveira, Valquíria Reis, Nayara Duarte, Barbara Malta; Hemominas – Juiz de For a (Minas Gerais) – Daniela Werneck; Hemominas; Montes Claros (Minas Gerais) – Rosimere Mota, José Wilson Sales, Maria Aparecida Souza, Rodrigo Ferreira; Fundação Hemope – Recife (Pernambuco) – Maria do Carmo Valgueir; Regina Gomes, Airly Goes Maciel, Rebeca Talamatu Dantas; Hemorio – (Rio de Janeiro) – Flavia Herculano, Ana Claudia Pereira, Ana Carla Alvarenga, Adriana Grilo, Fabiana Canedo; Instituto de Matemática e Estatística da Universidade de São Paulo - USP (São Paulo) – Marcio Moikawa, Mina Cintho Ozahata, Rodrigo Muller de Carvalho, João Eduardo Ferreira. US Investigators: RTI – Research Triangle Institute, International – Christopher McClure; National Institutes of Health, National Heart, Lung, and Blood Institute – Simone A. Glynn.

---

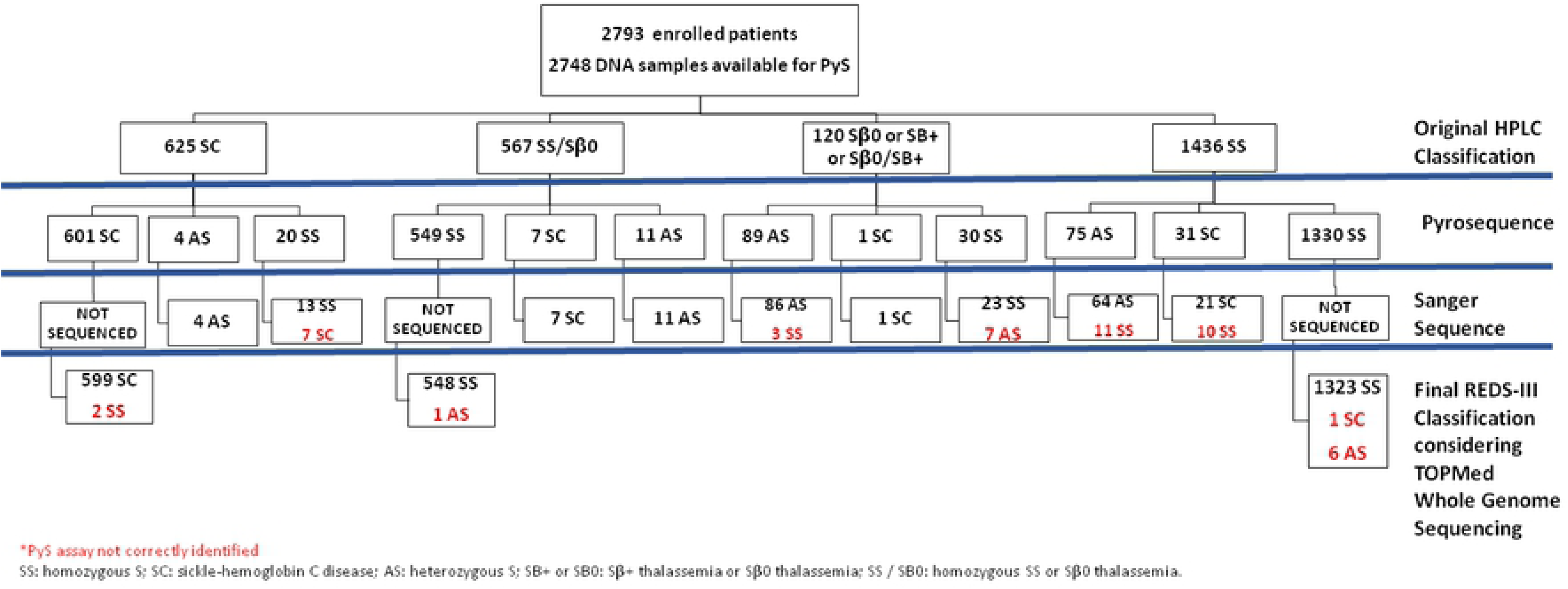

